# Characterisation of Quorum Sensing System and Its Role in Global Regulation in *Hafnia alvei*

**DOI:** 10.1101/812727

**Authors:** Jia Yi Tan, Wah-Seng See-Too, Peter Convey, Kok-Gan Chan

## Abstract

Quorum sensing (QS) is a regulatory process achieved via cell-to-cell communication that involves release and detection of autoinducers (AIs), and which occurs in a wide range of bacteria. To date, QS has been associated to events of pathogenesis, biofilm formation, and antibiotic resistance in clinical, industrial, and agricultural contexts. The main objective of this study was to characterise the role of *N*-Acyl homoserine lactone (AHL) type QS in *Hafnia alvei* FB1, a bacterial strain isolated from frozen vacuum-packed fish paste meatballs, via identification of QS core genes using a genomic approach, followed by comparative transcriptomic profiling between QS-deficient mutants and wild-type strains. H. alvei FB1 is known to produce two types of AHLs, namely, *N*-(3-oxohexanoyl) homoserine lactone (3OC6-HSL) and *N*-(3-oxooctanoyl) homoserine lactone (3OC8-HSL). The complete genome sequence of strain FB1 was obtained and a single gene for AHL synthase (*halI*) and its cognate receptor (*halR*) were identified. QS-deficient mutants of FB1 were constructed via the λ-Red recombineering method. Removal of the QS genes in strain FB1 affected mainly mechanisms in cell division and nutrient uptake, as well as resistance to a number of antibiotics, which are crucial for survival, adaptation and colonisation of both food and the host gut environment.

**Impact statement:** The *Hafnia* genus is known as opportunistic pathogen in both nosocomial and community-acquired infections, however, involvement and mechanism of pathogenesis of *Hafnia* in infection diseases is uncertain. We investigate the role of the signalling molecule, *N-*acyl homoserine lactones (AHLs), in a *Hafnia alvei* strain, since AHLs play important roles in pathogenicity, survival or adaptation in other pathogen. This comparative transciptomic study has revealed that AHLs are involved in mechanisms in cell division and nutrient uptake, as well as resistance to a number of antibiotics, which are crucial for survival, adaptation and colonisation of both food and the host gut environment. This finding provides insight and possible strategy to combat this opportunistic pathogen.

**Data summary:** Genome sequence is deposited in NCBI GenBank under accession number CP009706. The transcriptomic data, have been deposited in the Gene Expression Omnibus (GEO) database (https://www.ncbi.nlm.nih.gov/gds) under accession number GSE93000.

## Introduction

Quorum sensing (QS) is a form of cell-to-cell communication expressed by a wide range of bacterial species, achieved by the release and detection of signaling molecules called autoinducers (AIs) (1). *N-*acyl-homoserine lactone (AHL) is one such molecule, and is commonly produced by Proteobacteria. Individual cells monitor the changes in ‘quorum’ through detection of these small, often lipophilic, molecules in the environment and, in response, adjust the expression of a network of genes in a collective manner (2). AHL-based QS has been reported to be associated with various microbial activities of clinical, industrial, and agricultural importance, such as regulation on virulence expression (3-5), production of antibiotics (6, 7), biofilm formation (8-10), and food spoilage (11-13).

*Hafnia*, a bacterial genus characterised by rod-shaped, motile, flagellated, and facultatively anaerobic cells, belongs to the family *Hafniaceae. Hafnia alvei* has been identified to be among the enteric bacteria commonly involved in food spoilage (11, 12, 14, 15), and also as an opportunistic pathogen (16). The ability of *H. alvei* to survive at low temperature (15) has made it an interesting subject of study in controlling bacterial contamination in the food industry. Despite the common association of *H. alvei* with human gut microbiota (17-19), its status as a gut pathogen is still uncertain. Therefore, it is also important to confirm whether this common microbial contaminant of food could also be a source of gut infection.

The focus of this study, *H. alvei* FB1 is an AHL-producing strain recovered from vacuum-packed refrigerated fish paste meatballs (commonly known as ‘fish ball’). We have previously confirmed that the strain produces two oxo-AHLs with side chains of six or eight carbons in length (20).

In recent years, advancement in high-throughput sequencing and accessibility of powerful bioinformatics pipelines have enabled bacterial genomes and transcriptomes to be explored with increasing ease. Further, current DNA recombinant technology allows precise genome editing and provides a convenient tool in the study of QS regulons. This study investigated the genetic basis of AHL-based QS signalling in *H. alvei* FB1, as well as the regulatory role of QS in *H. alvei*, by studying the QS-deficient mutants in a comparative transcriptomics context.

## Methods

### Bacterial Strains, Plasmids and Culturing Conditions

All bacterial strains and plasmids used in this study are listed in Table 1 along with their relevant characteristics. All strains were cultured and maintained in Luria-Bertani (LB) medium. Unless stated otherwise, the growth temperatures for *Chromobacterium violaceum* CV026, *Escherichia coli* and *H. alvei* strains were 28°C, 37°C and 30°C, respectively.

**Table 1:**
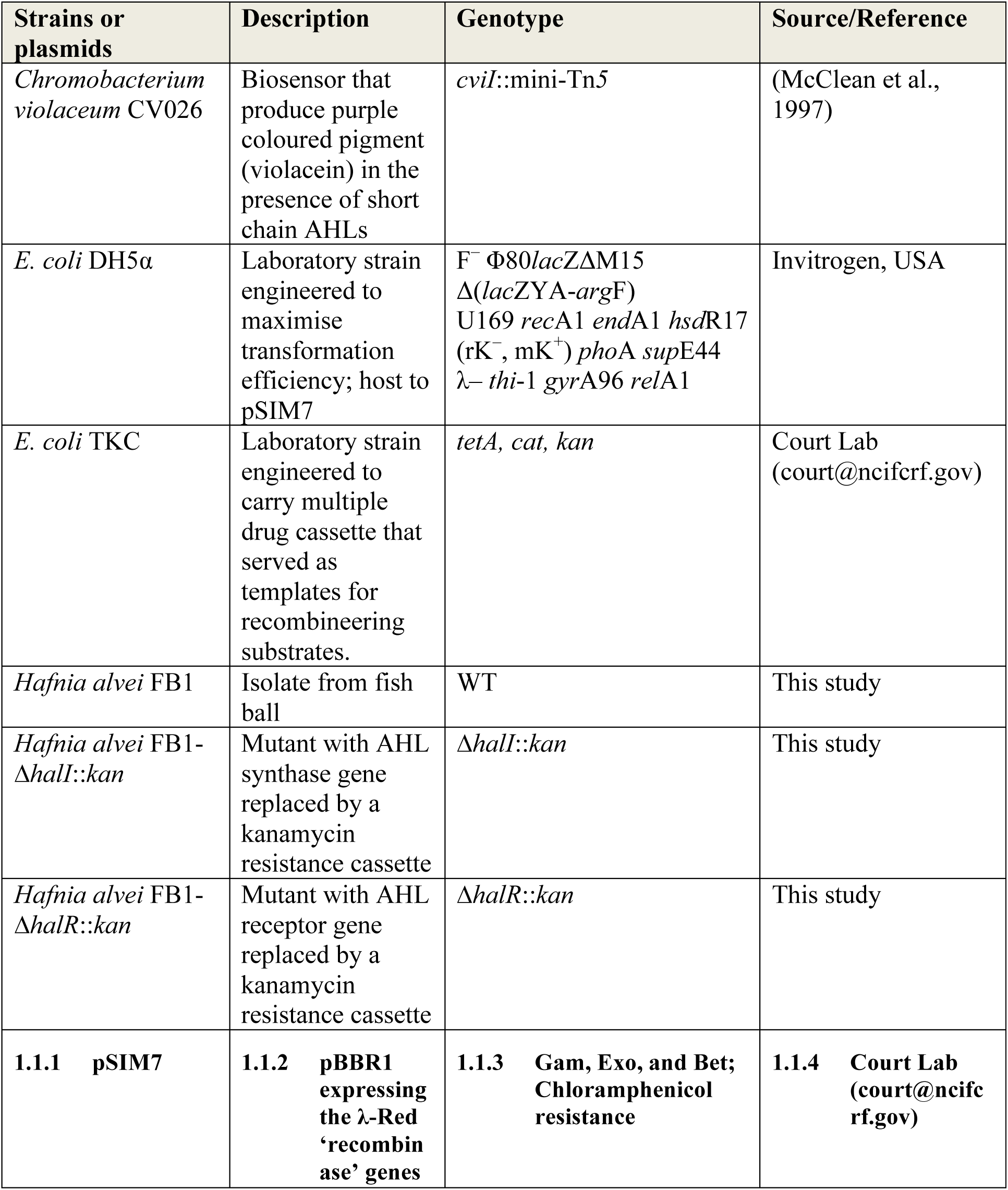
Bacterial strains and plasmids used in this study.

### Complete Genome Sequencing

The gDNA used in this study was extracted with the MasterPure™ DNA Purification Kit (Epicentre, USA) following the manufacturer’s protocol. The quality of gDNA extracted was assessed according to “PACBIO^®^ GUIDELINES FOR SUCCESSFUL SMRTbell™ LIBRARIES” prior to library preparation for sequencing. Preparation of SMRTbell template libraries was carried out with SMRTbell Template Prep Kit 1.0 (Pacific Biosciences, USA) following the manufacturer’s protocol for 20 kb template preparation. SMRT sequencing was performed using a PacBio RSII Sequencer (Pacific Biosciences, USA) with 3 SMRT cells.

Quality filtering and *de novo* assembly of the output reads were performed using the Hierarchical Genome Assembly Process (HGAP) pipeline version 3.0 on the Pacific Biosciences’ SMRT portal (21). Polished assemblies were generated from this module that incorporated Celera Assembler, BLASR mapper, and Quiver consensus caller algorithms. The assembled genome sequence was then annotated by SEED-based subsystem characterisation on the RAST server (22, 23) and Prokaryotic Genome Automatic Annotation Pipeline (PGAAP) (24).

### Identification of QS genes from the annotated data

Candidates of potential *luxI* and *luxR* homologues were identified from the annotated genome with the ‘BLAST search’ function on RAST. A search on the Conserved Domain Database (CDD) (25) was performed to identify the signature conserved domains on the candidate sequences. The sequences were aligned with other known homologues obtained from the GenBank database (https://www.ncbi.nlm.nih.gov/) and visualised using Jalview version 2. Phylogenetic analysis was performed using SeaView version 4 (26).

### Cloning and Expression of the *luxI* homologue

Identity and function of the *luxI* homologue were determined via cloning and expression study using Gene Synthesis service (GenScript, USA). The target DNA sequence was first synthesised in vector pUC57, and then sub-cloned into *E. coli* expression vector pGS21a by the service provider. The expression vector carrying the target sequence was transferred into expression host *E. coli* BL21 Star (DE3). The expression of the *luxI* homologue was examined *via* a cross-streak assay on the transformed host with a CV026 biosensor. AHLs produced were then identified on an LC-MS/MS platform as described previously (20).

### Construction of *halI* and *halR* replacement knockout mutants

Replacement knockout of QS genes in *H. alvei* FB1 was carried out according to Court Lab’s protocol “Recombineering: Using Drug Cassettes to Knock out Genes *in vivo*”, downloadable from the official website (https://redrecombineering.ncifcrf.gov/). Linear DNA substrates for recombineering were constructed through PCR amplification with 70-base hybrid primers (Table 2) consisting of 50 bases of homologous sequence that flanked the sequence to be replaced and 20 bases at the 3’ end that primed the amplification of the kanamycin resistance (KanR) cassette from the template strain (TKC) (27). The substrates were later introduced into heat-induced *H. alvei* FB1 cells harbouring pSIM7 *via* electroporation, in which the recombinases were actively expressed. The recovered cells were then plated on a kanamycin plate (35 µg/mL) to select for mutants that carried the KanR cassette in the place of QS genes. The results of recombineering were then confirmed by colony PCR using primers (*halI*_u-F, *halI*_d-R, *halR*_u-F, and *halR*_d-R with Screen_KanR-F and Screen_KanR-R, as described in Table 2) that amplified the junctions where recombination took place. Cross-streaking with CV026 was also performed on the *H. alvei* FB1Δ*halI*::KanR to confirm that the AHL synthase function had been successfully removed.

**Table 2:**
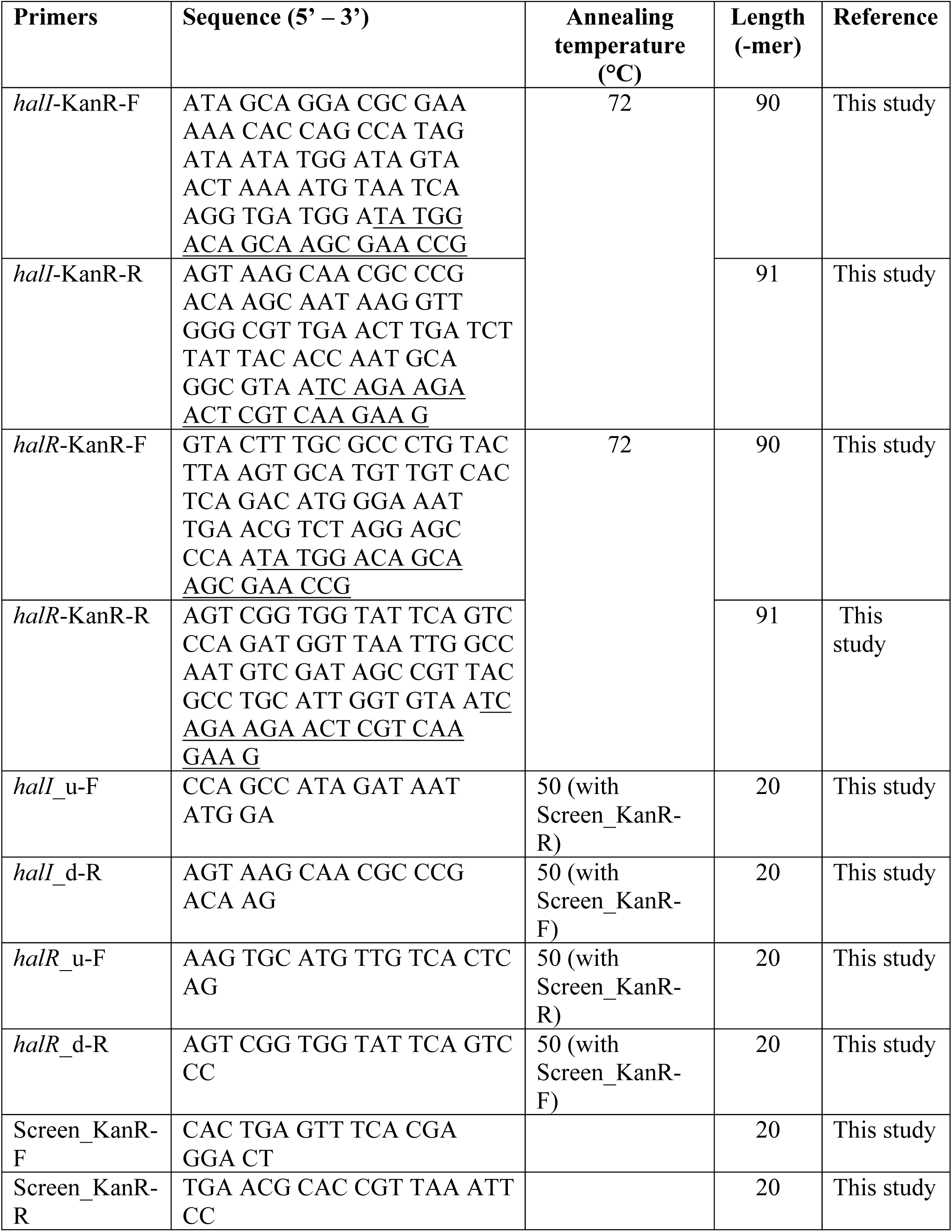
Oligonucleotides used in PCR amplification for mutant construction.

### RNA-Seq

*H. alvei* FB1 WT and mutant cells were grown in 200 mL LB broth at 30°C with shaking at 220 rpm. Readings of optical density (OD_600_) were taken every hour. The total RNA was stabilised immediately using Qiagen RNAprotect bacterial reagent (Qiagen, USA) when the cells were harvested at the desired OD_600_ (0.5 and 1.5), following the manufacturer’s protocol. Total bacterial RNA was purified using the Qiagen RNeasy kit (Qiagen, USA) following the manufacturer’s protocol. Purified RNA samples were eluted in 100 µL sterile RNase-free water. Total RNA from three biological replicates was obtained from independent cultures to evaluate the reproducibility of RNA-Seq data. Purity of RNA samples was assessed using a Nanodrop spectrophotometer (Thermo Scientific, USA), and RNA quantification was carried out using the Qubit^®^ RNA HS Assay Kit (Invitrogen, USA). The quality of the extracted RNA samples was assessed with an Agilent RNA 6000 Nano Kit (Agilent Technologies, USA) using a 2100 Bioanalyzer.

RNA samples with RNA Integrity Number (RIN) >7.0 were chosen to proceed to rRNA depletion using Ribo-Zero™ rRNA Removal Kits (Bacteria) (Epicentre, USA) prior to cDNA synthesis. The quality of the rRNA-depleted RNA was assessed with an Agilent RNA 6000 Pico Chip (Agilent Technologies, USA) using a 2100 Bioanalyzer. Library preparation was performed following the manufacturer’s protocol using the Illumina ScriptSeq™ v2 RNA-Seq Library Preparation Kit (Epicentre, USA). Quality of the RNA-Seq transcriptome library was examined using Agilent 2100 High Sensitivity DNA Kit. Quantification of the library was assessed using the Qubit^®^ dsDNA HS Assay Kit (Life Technologies, USA) and qPCR (KAPA Biosystems) before being subjected to normalisation. The normalised samples (4 nM) were denatured with 0.2 N NaOH and diluted 20 pM using pre-chilled Hybridisation Buffer (HT1) (Illumina, USA). The 20 pM transcriptome libraries were further diluted to 10 pM with pre-chilled HT1 buffer prior to whole transcriptome sequencing on a MiSeq platform.

RNA-seq read quality assessment was carried out using FastQC (version 0.11.2) (28). Rockhopper (29, 30) was used to align the paired end reads against the *H. alvei* FB1 reference genome (GenBank accession number CP009706). Differential expression analysis was then performed with the DESeq2 package using the raw counts generated from Rockhopper as inputs (*p*_adjust_ value/*q* < 0.01) (31). Gene Ontology (GO) and pathway enrichment analyses of differentially expressed genes (DEGs) were performed using The Database for Annotation, Visualization and Integrated Discovery (DAVID) v6.8 (32, 33)

### qPCR

Prior to qPCR, total RNA was converted to cDNA using the QuantiTect^®^ Reverse Transcription Kit (Qiagen, Germany) following manufacturer’s protocol. qPCR was performed with a CFX96 Touch™ Real-Time PCR Detection System (Bio-Rad, USA) using the primers listed in Table 2. Expression levels in ΔΔCq of each gene were determined from analyses using Bio-Rad CFX Manager version 3.1 (Bio-Rad, USA).

## Results

### Identification of QS genes of *H. alvei*

The genome sequence of *H. alvei* FB1 is deposited in GenBank under the accession number CP009706. A homology search using LuxI and LuxR amino acid sequences via the ‘BLAST search’ function on the RAST server revealed the presence of a pair of open reading frames (ORF) at position 2,668,370 – 2,669,738 in the genome that contained the signature conserved domains of LuxI and LuxR homologues. The pair consisted of a 648-bp *halI* (Locus tag AT03_RS12360) along with a 741-bp *halR* (AT03_RS12355). The pair was in convergent orientation, with a 20-bp overlap at the 3’-ends (Figure 1A).

**Figure 1:**
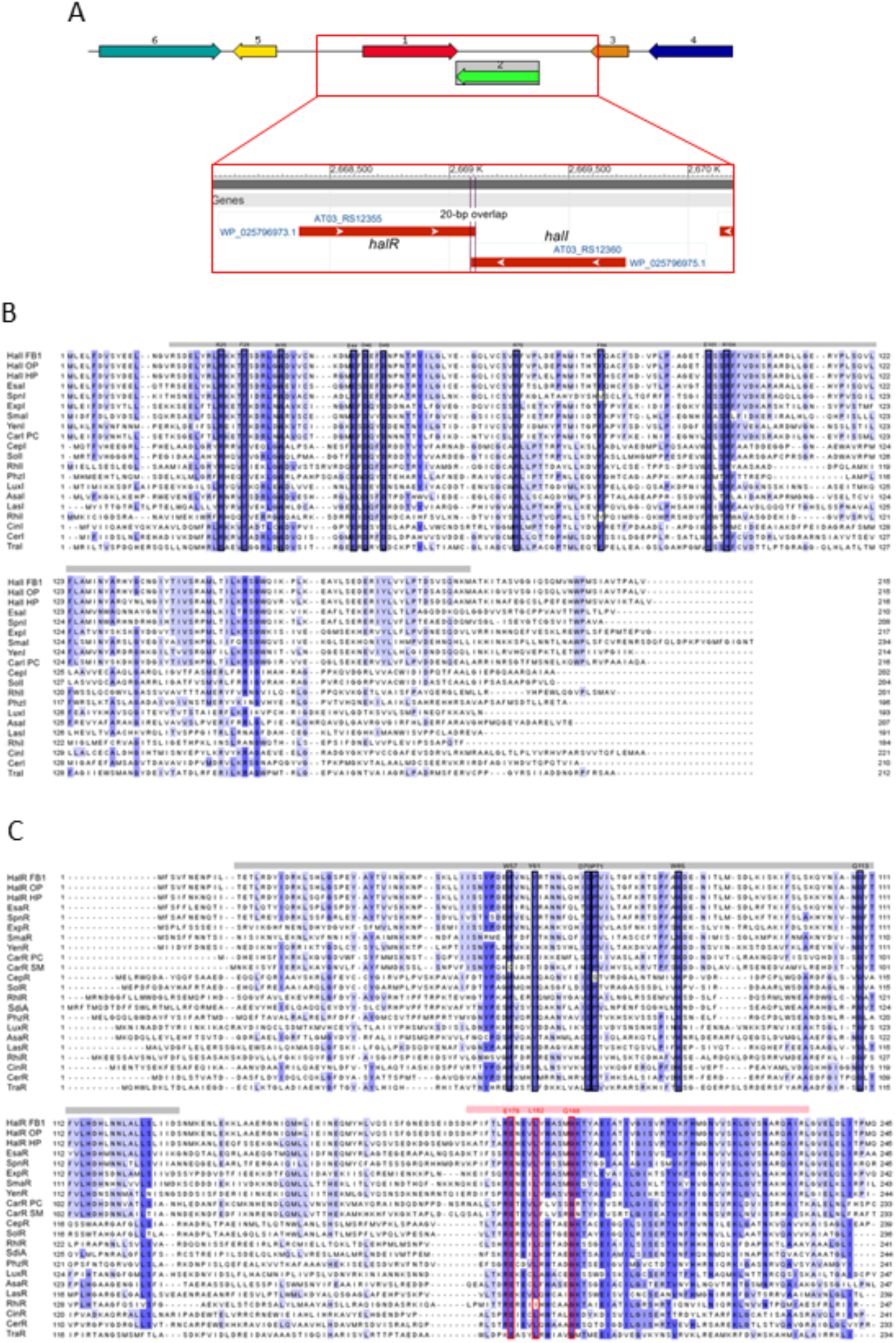
Physical map and sequence analyses of HalI and HalR in *H. alvei* FB1. (A) The arrows indicate the coding regions of *halI* and *halR*, along with the orientation and the flanking genes. 1. *halR*; 2. *halI*; 3. Hypothetical protein; 4. Probable secreted protein; 5. Permease of the drug/metabolite transporter (DMT) superfamily; 6. *umuC*. (B) MSA of amino acid sequences of HalI and other known LuxI homologues. Ten key conserved residues according to their positions in TraI are highlighted. OP: *O. proteus*; HP: *H. paralvei*; PC: *P. carotovorum*. (C) MSA of amino acid sequences of HalR and other known LuxR homologues. Nine key conserved residues within DNA binding domain according to their positions in TraR are highlighted. OP: *O. proteus*; HP: *H. paralvei*; PC: *P. carotovorum*.

Multiple sequence alignment (MSA) was performed between HalI amino acid sequence and other known homologues of LuxI (Figure 1B). Ten conserved key amino acid residues, namely R25, F29, W35, E44, D46, D49, R70, F84, E101 and R104 according to their positions in TraI as reported by Fuqua and Greenberg (34), were identified in the sequence of HalI. Likewise, MSA on HalR and other known homologues of LuxR revealed the presence of six conserved key residues (W57, Y61, D70, P71, W85 and G113 according to their positions in TraR) within the autoinducer binding domain and three (E178, L182, and G188) within the DNA binding domain (Figure 1C), as reported by Subramoni et al. (35).

Phylogenetic trees were built to illustrate the phylogenetic relationships between the HalI and HalR proteins and other LuxI and LuxR homologues of known functions (Figures 1, 2). Both trees shared roughly similar topologies, consisting of two major clusters. The top clusters consisted solely of members of γ-Proteobacteria, whereas the bottom ones contain those of α-, β- and γ-Proteobacteria. All sequences from *Hafnia* spp. formed a single clade in both trees, except for an orphan LuxI homologue (YspR) in *H. paralvei* bta3-1 that clustered with the more distantly related SmaR of *Serratia* sp. ATCC 39006. Both HalI and HalR were more closely related to YspI-YspR in *Yersinia pestis* and YtbI-YtbR in *Y. pseudotuberculosis*, compared with *Edwardsiella tarda* ATCC 15947^T^ that also belongs to the *Hafniaceae*. The LuxR tree clearly shows that all the HalR sequences formed a monophyletic clade with the sequences of LuxR homologues that are known to establish quorum-hindered activities.

**Figure 2:**
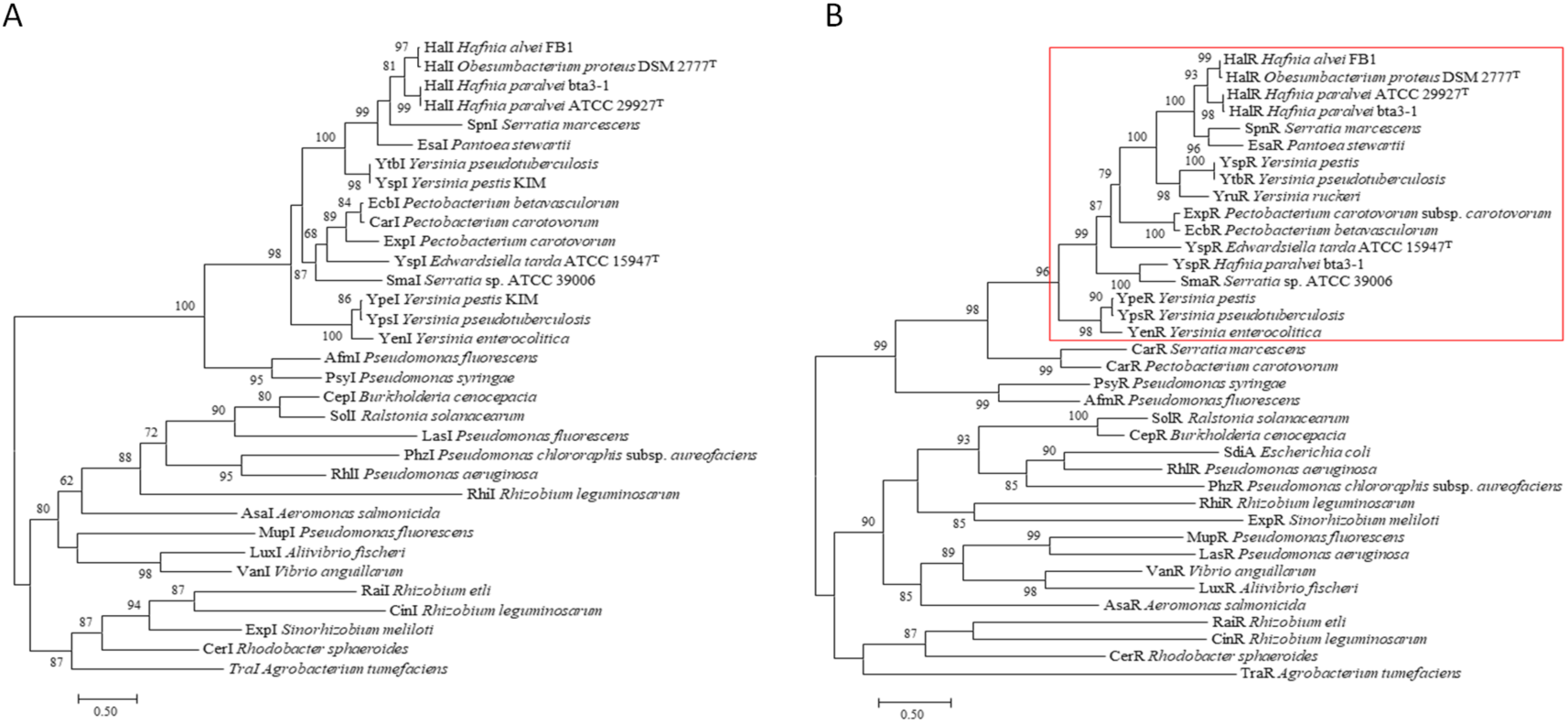
(A) Phylogenetic tree showing the evolutionary distance of HalI of *H. alvei* FB1 and other LuxI homologues inferred using the Maximum Likelihood (PhyML) method based on the LG matrix-based model. The tree with the highest log likelihood (−11301.6) is shown. The aLRT branch support values are shown next to the branches. The tree is drawn to scale, with branch lengths measured in 0.5 substitutions per site (bar). (B) Phylogenetic tree showing the evolutionary distance of HalR of *H. alvei* FB1 and other LuxR homologues inferred using the Maximum Likelihood (PhyML) method based on the LG matrix-based model. The tree with the highest log likelihood (−14799.2) is shown. The aLRT branch support values are shown next to the branches. The tree is drawn to scale, with branch lengths measured in 0.5 substitutions per site (bar). The monophyletic clade that contained HalR and other LuxR homologues known to establish quorum-hindered properties is highlighted in the red box.

### Expression of *halI* in a Foreign Host

To prove the functionality of the identified AHL synthase, the *halI* sequence was cloned into a pGS-21a vector and expressed in host *E. coli* BL21 Star (DE3). Cross-streaking of the transformed BL21 Star (DE3) induced purple pigmentation of the CV026 biosensor (Figure S1). The same AHL profile produced by *H. alvei* FB1 (3OC_6_-HSL and 3OC_8_-HSL) was detected using LC-MS/MS (Figure S2).

### Mutant Construction *via* λ-Red Recombineering

Two mutants, namely, Δ*halI*::kanR and Δ*halR*::kanR were constructed using the λ Red recombineering technique. In each mutant, either *halI* or *halR* was replaced with a kanamycin resistance cassette. Verification of the mutants was performed via PCR on colonies picked from LBA plates containing 35 mg/mL kanamycin. The presence of amplification products encompassing the junctions between the flanking regions and a stretch of the intragenic region of KanR cassette indicated the success of the replacement knockout. The PCR products were sent for Sanger sequencing to confirm the sequences (Figure S3). The Δ*halI*::kanR mutants, when tested by cross-streaking with biosensor CV026, were unable to induce purple pigment production (Figure S4).

### RNA-Seq

To examine the pattern of global gene expression affected by HalR regulon, RNA-Seq was performed for the two mutants using strain FB1 WT as a control. Transcriptomes of each strain were acquired at early- and mid-exponential phase (OD_600_ = 0.5 and 1.5, respectively). Three libraries of FB1 WT, Δ*halI*::KanR, and Δ*halR*::KanR were generated for each growth phase, which yielded total reads ranging from 2,124,060 to 4,957,038. The Rockhopper software was able to map between 504,685 and 3,344,049 reads to the reference genome. The sequencing output, along with the raw counts, have been deposited in the Gene Expression Omnibus (GEO) database (https://www.ncbi.nlm.nih.gov/gds) under accession number GSE93000.

### Quality Assessment of the Replicates

The principal component analysis (PCA) plot based on the raw counts showed that, for the samples harvested at OD_600_ 0.5, the three replicates of Δ*halR*::KanR samples were clearly separated from the WT and Δ*halI*::KanR samples along the PC1 axis, which contributed 94.662% of the variation, while the latter two clustered close to each other on this axis (Figure 3A). For the samples harvested at OD_600_ 1.5, the replicates formed distinct clusters in different quadrants of the plot, with the Δ*halI*::KanR mutant forming a cluster more distinct from the WT samples along the PC1 axis (which explained 79.838% of variation) than the Δ*halR*::KanR samples (Figure 3B).

**Figure 3:**
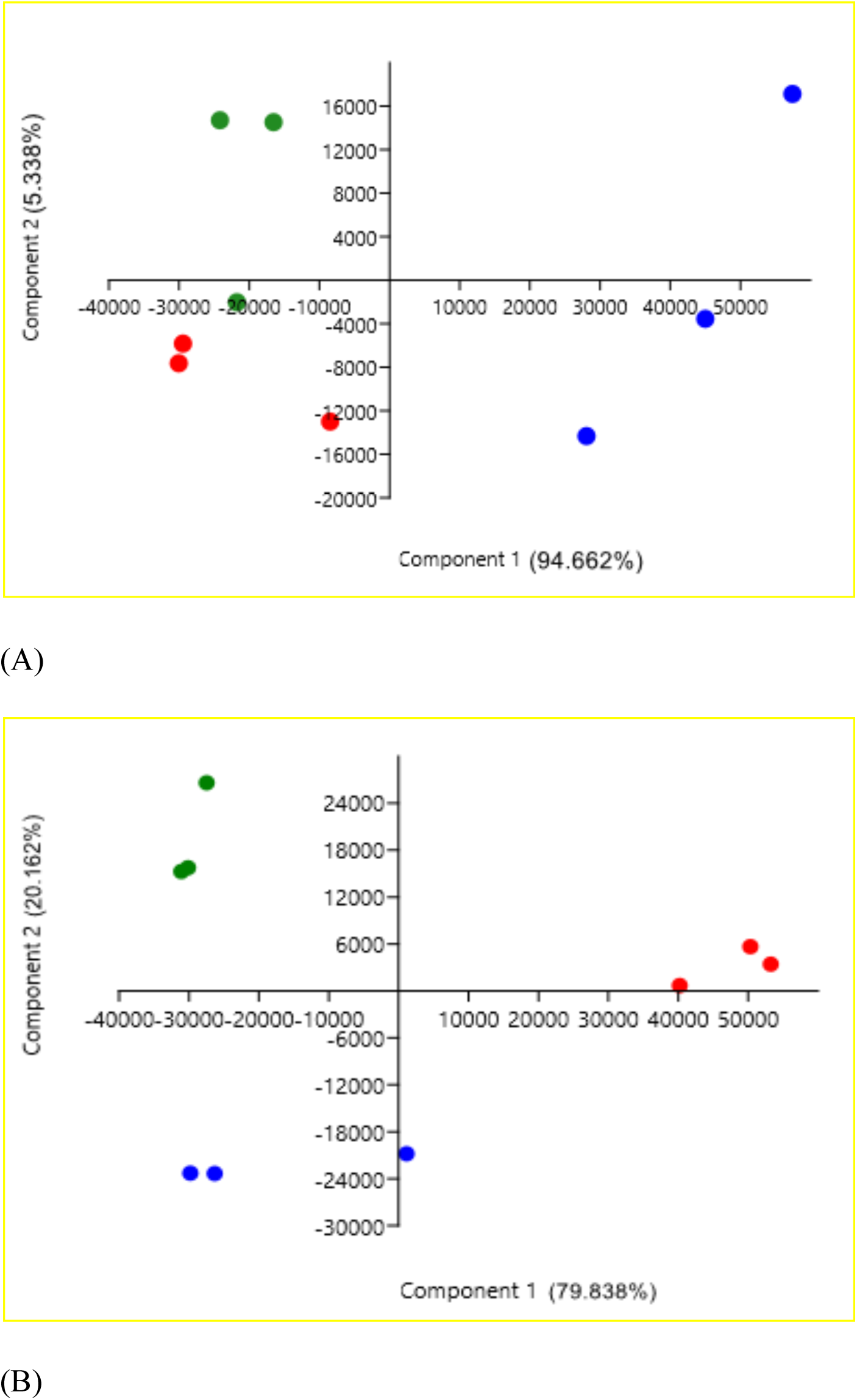
PCA scatter plots showing the variation between samples and consistency between replicates. Triplicates of *H. alvei* FB1 WT, Δ*halI*::KanR, and Δ*halR*::KanR are represented as green, red, and blue dots, respectively. (A) Samples harvested in early exponential phase. (B) Samples harvested in mid-exponential phase.

### Transcriptomic Analysis of QS Mutants

No significant differential expression (log_2_ fold change [FC] > 1 or < −1) was observed in the Δ*halI*::KanR mutant at a *q*-value of 0.01, aside from *halI*. A total of 687 genes were found to be differentially expressed in the Δ*halR*::KanR mutant, of which 511 were up-regulated while 176 were down-regulated. The distribution of DEGs is illustrated as MA plots in Figure 4A, B.

**Figure 4:**
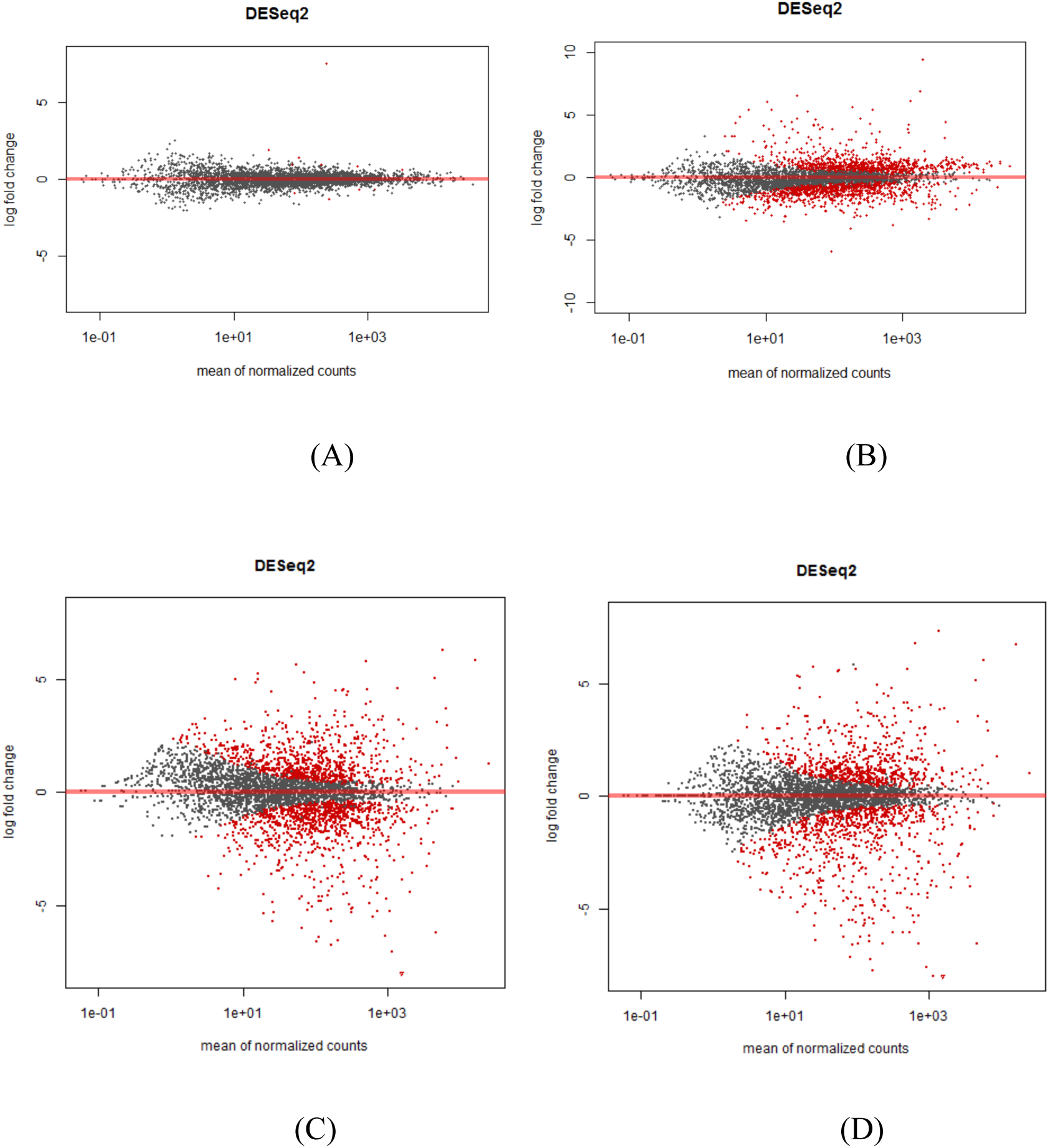
MA plots showing the distribution of DEGs in (A) Δ*halI*::KanR and (B) Δ*halR*::KanR against WT in early exponential phase and (C) Δ*halI*::KanR and (D) Δ*halR*::KanR against WT in mid-exponential phase. Grey dots: genes with log_2_ FCs below 1 and above −1; red dots: genes with log_2_ FCs above 1 and below −1.

A total of 825 and 740 genes were differentially expressed in the Δ*halI*::KanR and Δ*halR*::KanR mutants, respectively. In the former, 392 were up-regulated while 433 were down-regulated, while in the latter, 371 showed positive fold-changes against the WT strain while 369 showed negative. The distribution of DEGs is illustrated as MA plots in Figure 4C, D, and the overlapping genes between both mutants are shown in Venn diagrams (Figure 5) FC of genes in each experimental condition are listed in Table S1.

**Figure 5:**
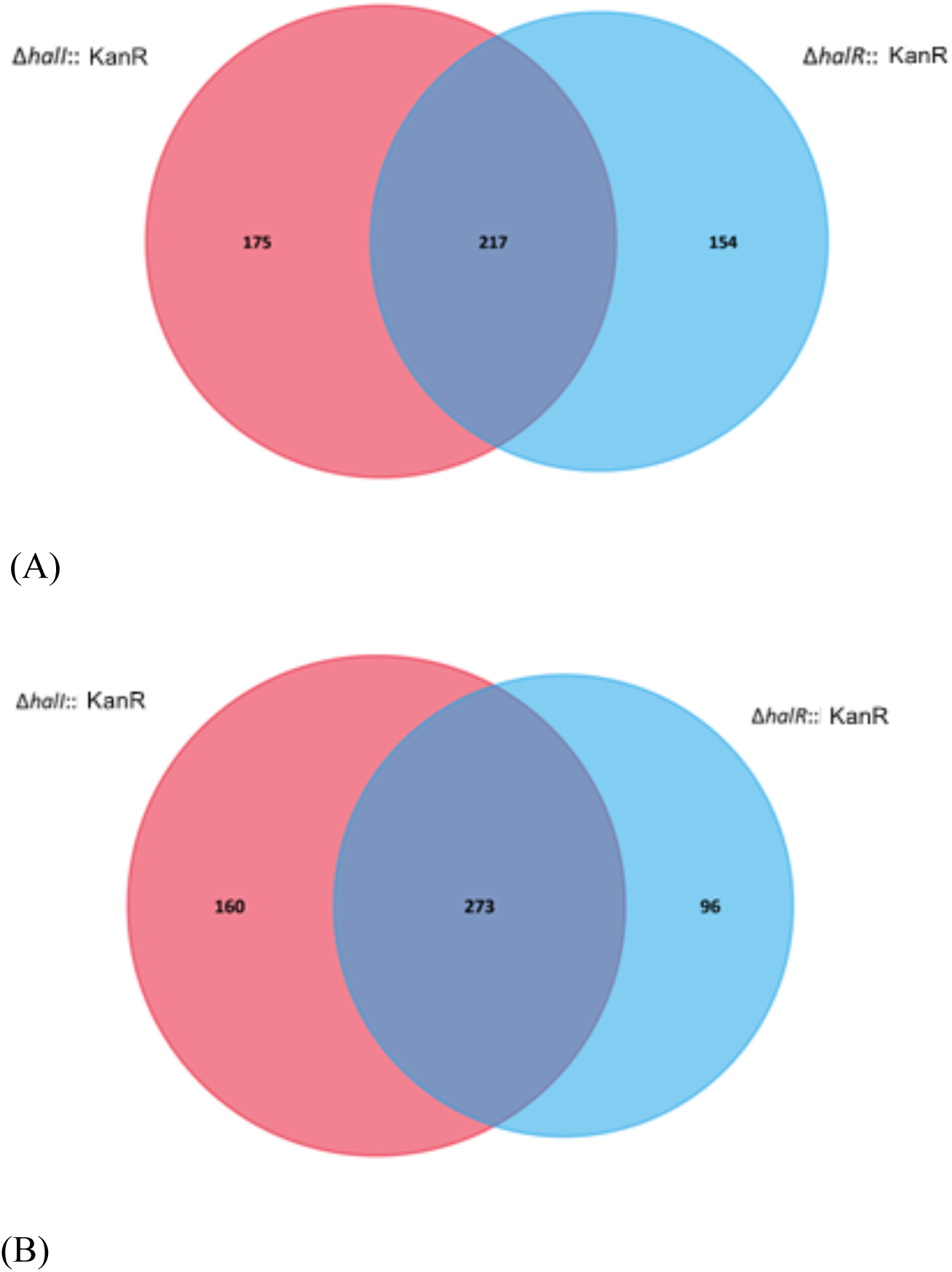
Venn diagrams showing the overlapping DEGs between Δ*halI*::KanR and Δ*halR*::KanR mutants in mid-exponential phase. (A) A total of 217 up-regulated genes (log_2_ FC > 1, *q-*value < 0.01) are shared between Δ*halI*::KanR (red circle) and Δ*halR*::KanR (blue circle), which contain 175 and 154 unique up-regulated genes, respectively, (B) A total of 273 down-regulated genes (log_2_ FC < −1, *q*-value < 0.01) are shared between Δ*halI*::KanR (red circle) and Δ*halR*::KanR (blue circle), which contain 160 and 96 unique down-regulated genes, respectively.

For the Δ*halR*::Kan mutant in exponential phase, both up- and down-regulated gene lists were topped by genes involved in carbohydrate transport and metabolism. Up-regulated genes were mainly involved with ribose metabolism, whereas down-regulated genes involved in mannitol, maltose and galactose uptake were among those with the highest FCs. In mid-exponential phase, genes involved in amino acid transport and metabolism were most abundant in the lists of both mutants. Genes involved with maltose transport, for instance maltoporin, which were down-regulated in the early exponential phase, were up-regulated in the mid-exponential phase.

### GO and Pathway Enrichment Analysis of QS-dependent Genes

GO enrichment analysis was performed on the DEGs to gain an insight into the QS-regulated genes. At the early exponential phase, out of the 511 up-regulated genes in the Δ*halR*::KanR mutant, 438 were assigned to 197 GO terms, of which five terms were enriched (*q*-value < 0.01). Of the 176 down-regulated genes, 114 were assigned to 58 terms, of which only two were enriched (Figure 6). The GO terms enriched in the up-regulated DEGs included fatty acid biosynthetic processes, cytoplasm, ATP binding, RNA binding, and tRNA binding, whereas the down-regulated genes included those involved in the phosphoenolpyruvate-dependent sugar phosphotransferase system and in response to stress.

**Figure 6:**
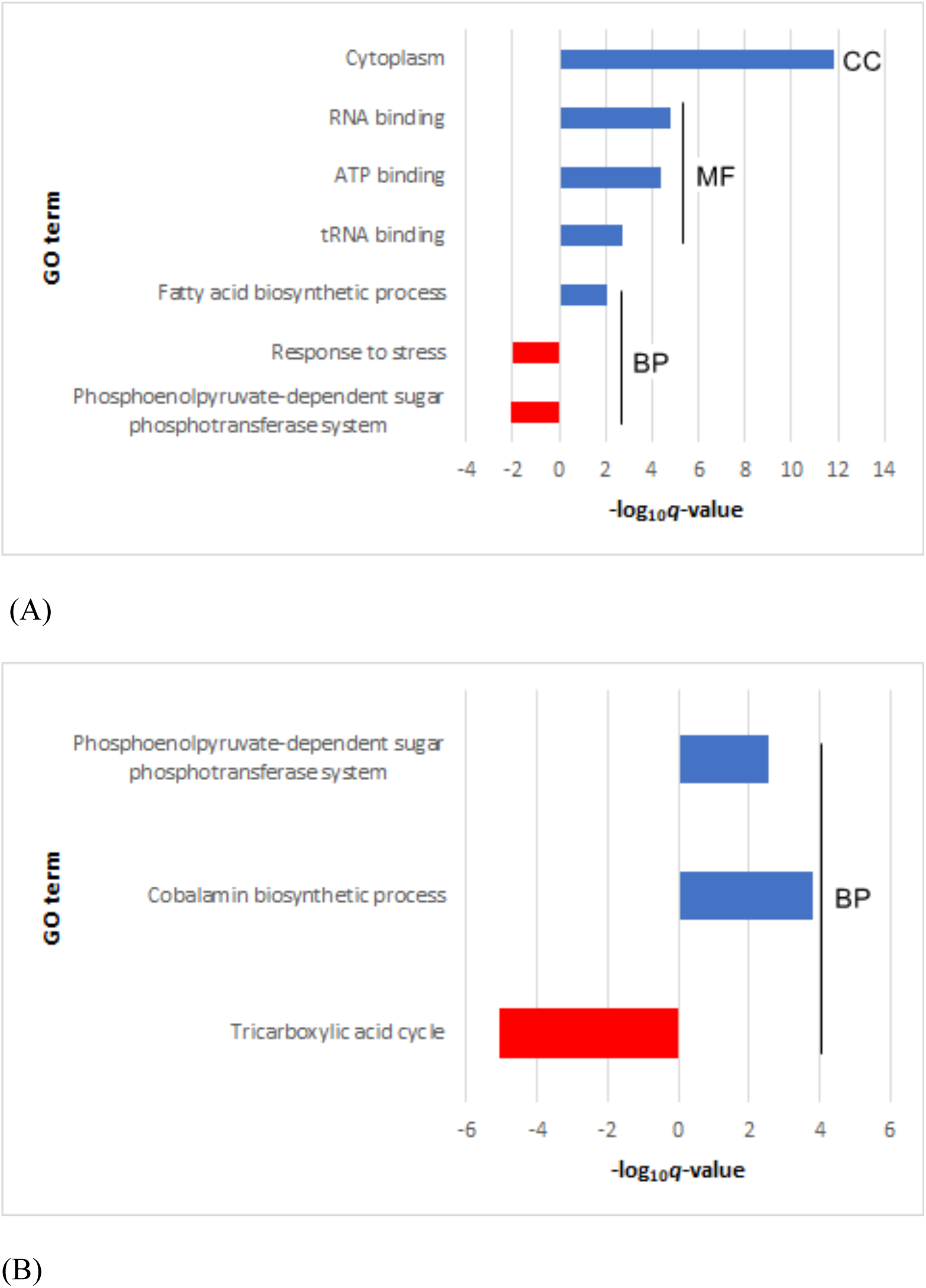
GO enrichment grouping of QS-dependent genes in *halR*::KanR in (A) early exponential phase and (B) mid-exponential phase. CC: Cellular content; MF: Molecular function; BP: Biological process.

In the mid-exponential phase, 304 out of 371 up-regulated genes in the Δ*halR*::KanR mutant were assigned to 180 GO terms, and two (phosphoenolpyruvate-dependent sugar phosphotransferase system and cobalamin biosynthesis process) were significantly enriched. Of the 263 GO terms accounting for 317 of the 369 down-regulated genes, only one (TCA cycle) was significantly enriched. In the *halI* mutant, genes with positive and negative FCs were assigned to 132 and 138 GO terms, respectively. None of these terms were enriched at *q*-value = 0.01.

DEGs were also mapped into the KEGG pathway database to identify pathways that were significantly enriched in each experimental condition (Figure 10). In early exponential phase, the up-regulated genes in the Δ*halR*::KanR mutant were assigned to a total of 80 pathways, of which 12 were significantly enriched at the cut-off of *q*-value = 0.01. These pathways included primarily those involved in biosynthesis of molecules and functions that involved nucleotides. Six of the 59 pathways to which the down-regulated genes in the Δ*halR*::KanR mutant were assigned were significantly enriched. As in the GO ontology, the phosphotransferase system (PTS) was involved, along with several other pathways involved in carbohydrate metabolism.

In mid-exponential phase, the number of down-regulated pathways in the Δ*halR*::KanR mutant far exceeded those that were up-regulated. A total of 30 out of 80 pathways were significantly enriched. These pathways were mostly involved in amino acid metabolism. The Δ*halI*::KanR mutant in the same growth phase showed a much smaller number of enriched pathways than did the Δ*halR*::KanR, but they were still dominated by down-regulated genes. Most of these pathways overlapped those affected in the *halR* counterpart, except for beta-lactam resistance.

### Validation of RNA-Seq Data by qPCR

To validate the RNA-Seq results, six genes were randomly chosen for qPCR. The log_2_ FCs of qPCR were calculated and plotted against that of RNA-Seq (Figure 8). The correlation coefficients were all above 0.99, indicating that the results obtained were reliable

**Figure 7:**
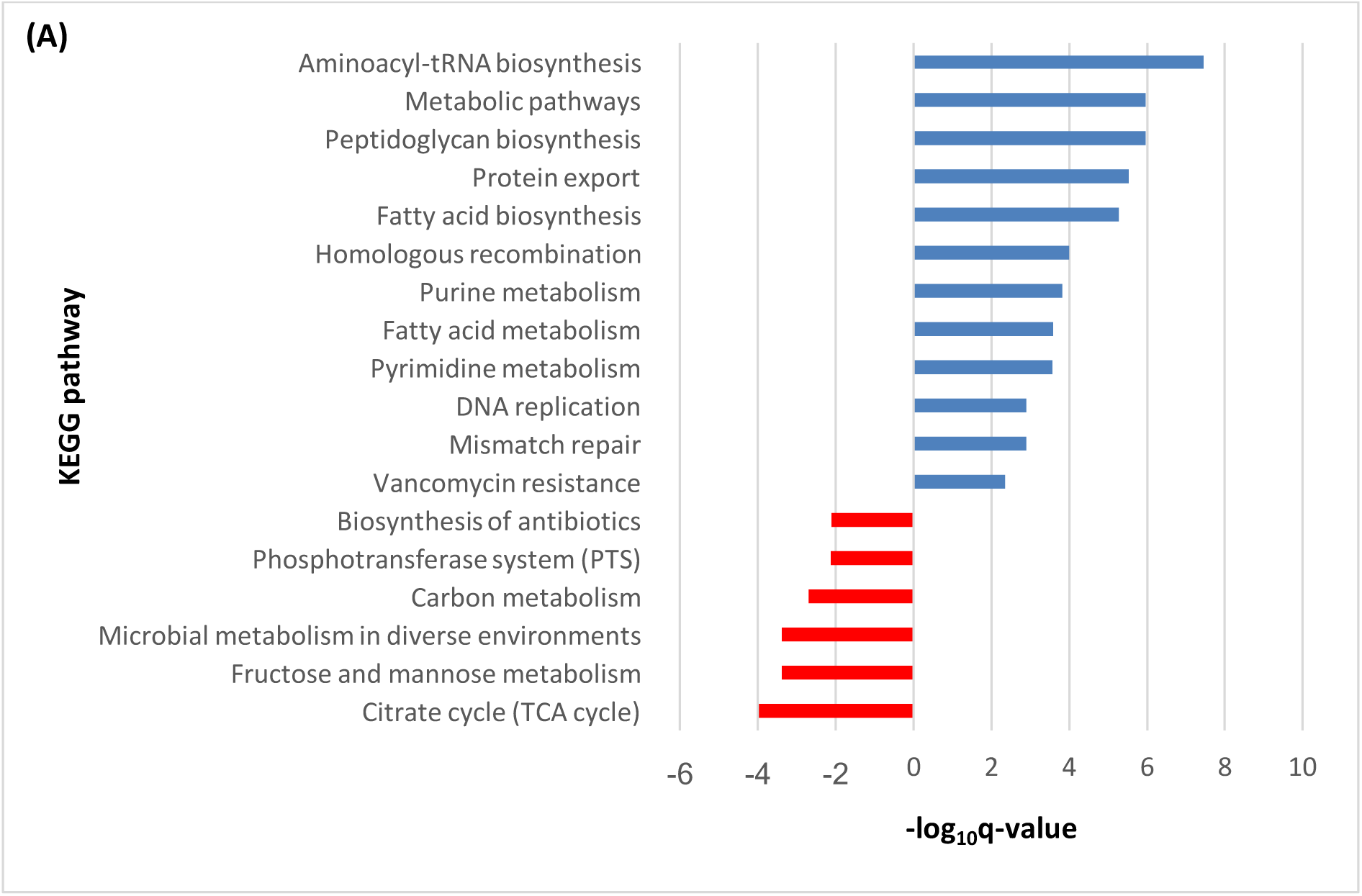
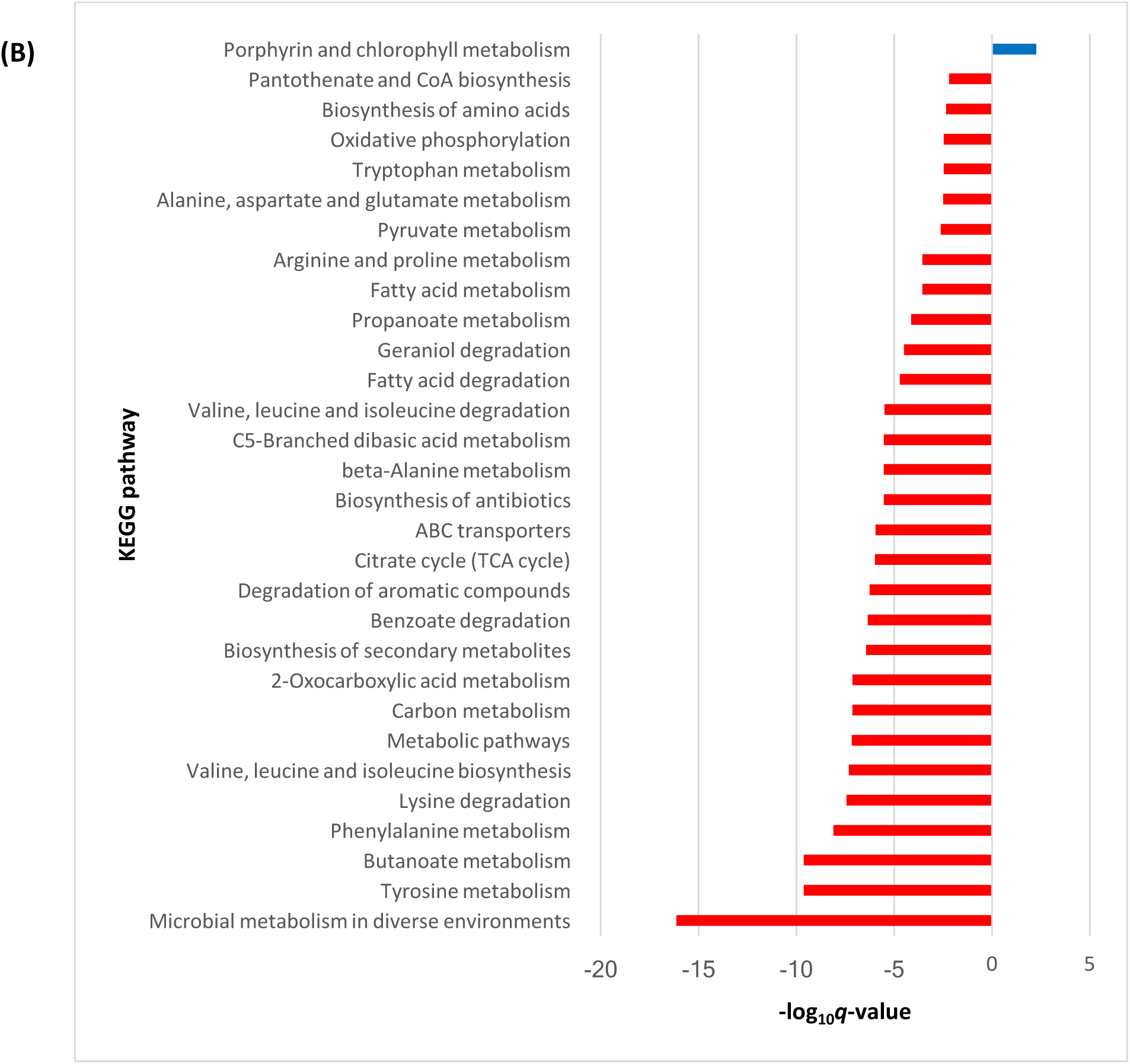
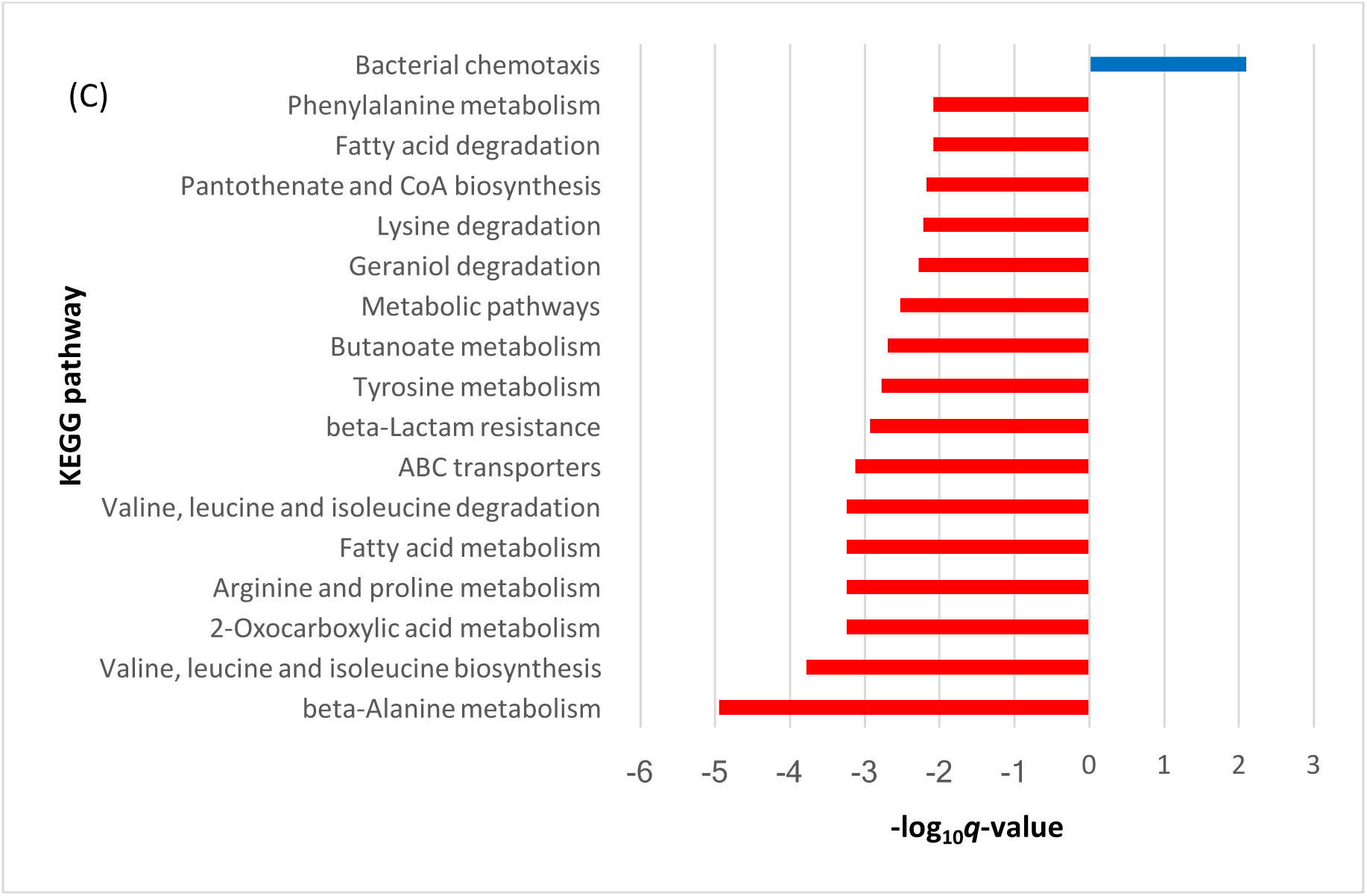
KEGG pathways enriched in (A) Δ*halR*::KanR mutant in early exponential phase, (B) Δ*halR*::KanR mutant in mid-exponential phase, and (C) Δ*halI*::KanR mutant in mid-exponential phase.

**Figure 8:**
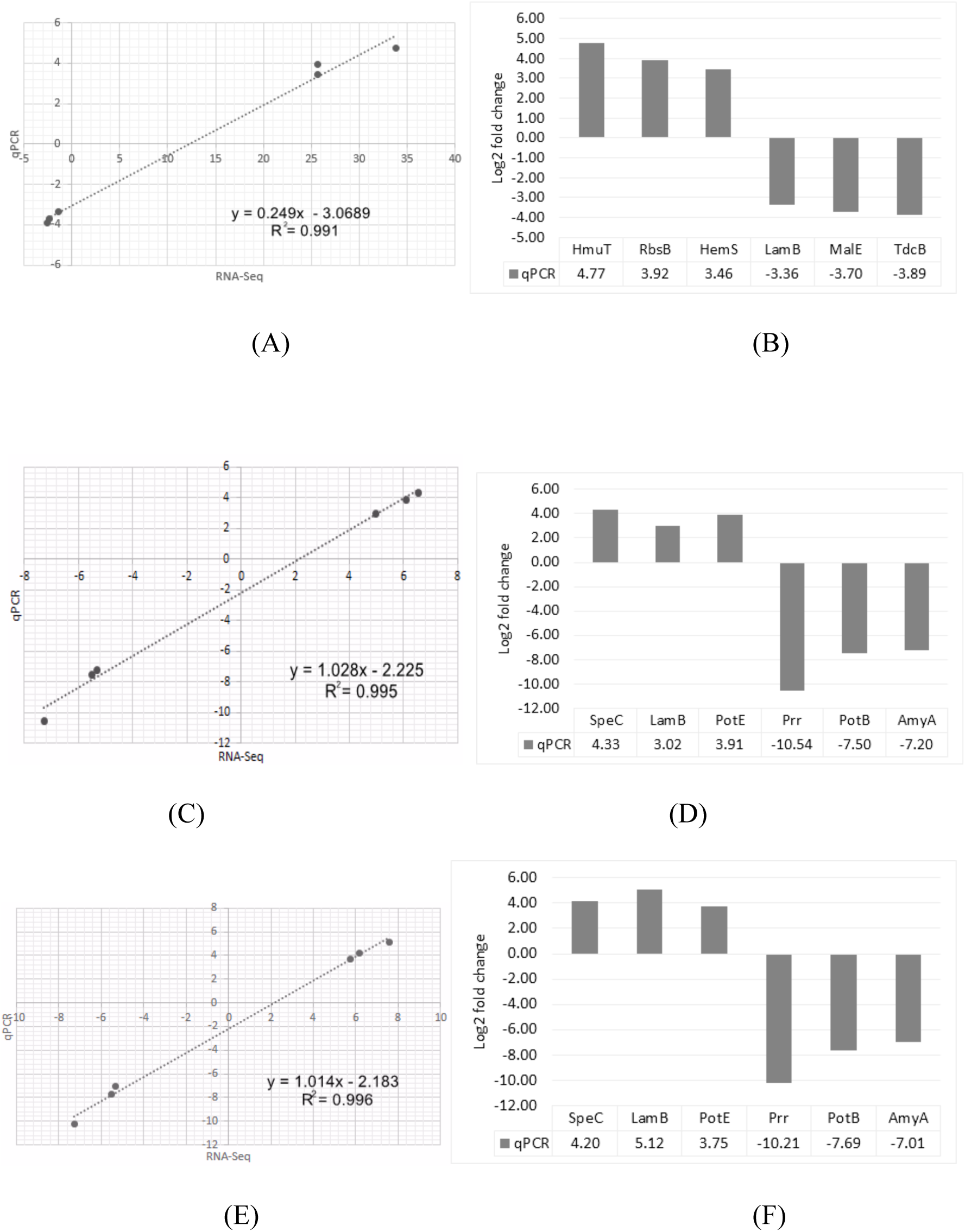
qPCR validation of RNA-Seq results. Each log_2_ ratio of FCs calculated from qPCR was compared with that of the RNA-seq data. (A) Correlation of the FCs between RNA-seq (x-axis) and qPCR (y-axis) of Δ*halR*::KanR in the early exponential phase. (B) Log_2_ FCs of Δ*halR*::KanR in the early exponential phase. (C) Correlation of FCs between RNA-seq (x-axis) and qPCR (y-axis) of Δ*halR*::KanR in the mid-exponential phase. (D) Log_2_ FCs of Δ*halR*::KanR in the mid-exponential phase. (E) Correlation of fold changes between RNA-seq (x-axis) and qPCR (y-axis) of Δ*halI*::KanR in the mid-exponential phase. (F) Log_2_ FCs of Δ*halI*::KanR in the mid-exponential phase.

## Discussion

Having examined *in silico* the characteristics of the candidates of QS genes in *H. alvei* FB1, the strain, unlike some other members of Proteobacteria, possessed only a single pair of *luxI-luxR* homologues. The presence of conserved domains and amino acid sites that are signature to the group of proteins, along with the orientation of the genes, provide strong evidence supporting the identity of the QS gene candidates. Phylogenetic trees constructed using choices of sequences included in the analyses adapted from a review by Tsai and Winans (36) revealed that the amino acid sequences of HalR proteins obtained from members of the genus *Hafnia* fell into a monophyletic clade with other EsaR-like proteins. This, along with other traits, such as a convergent orientation with overlapping 3’-ends and responding to 3OC_6_-HSL, suggested the possibility that HalR might establish similar quorum-hindered behaviour. An intermingling pattern between sequences originating from genomes that belonged to different classes of Proteobacteria was observed in one of the major clusters in both phylogenetic trees. This observation, along with the distant phylogenetic relationship between the orphan LuxR homologue with HalR, is likely to reflect the occurrence of HGT in the past, as has been reported in previous studies (37, 38).

According to the results of the transcriptomic study, Δ*halI*::KanR and Δ*halR*::KanR mutants showed difference in the functional distribution patterns of DEGs, which supports the proposition that HalR proteins belong to the quorum-hindered group. At early exponential phase, no significant DEG was observed in Δ*halI*::KanR, whereas a large number of DEGs were found in Δ*halR*::KanR. We suggest that, at OD_600_ of 0.5, the concentration of AHLs within the culture environment was still too low to affect the expression of any genes. Therefore, the mutant lacking AHL-synthase established a gene expression profile similar to the WT strain. The observation that removal of the *halR* gene resulted in a very different expression profile pointed to the possibility that there were HalR proteins already binding at the promoter regions of certain genes at this early stage, playing a regulatory role in the absence of sufficient QS signals. A much larger number of up-regulated genes was observed in the Δ*halR*::KanR mutant compared with those that were down-regulated, indicating that the major role of the HalR protein at this stage could be repression of certain functions in an early growth phase.

Later, in the mid-exponential phase, the gene expression profiles in the two mutants changed. As illustrated by PCA, the Δ*halR*::KanR mutant now clustered closer to the WT strain, implying that, with the increasing concentration of AHL molecules over time, the HalR proteins in FB1 WT were then released from their binding sites and the expression levels of the previously repressed genes became similar to the unrepressed ones in Δ*halR*::KanR. At this stage, DEGs could also be observed in the Δ*halI*::KanR mutant. More than half of these genes showed changes in the same directions in both mutants, which reflected a scenario more complex than classical examples of ‘quorum-hindered apo-proteins’, in which the changes in both mutants were expected to be in opposite directions. This observation suggests that a more complicated regulatory mechanism could be involved in the QS system in *H. alvei*.

### Functional Distribution of QS-regulated Genes

The enrichment analyses revealed that, in the early exponential phase, a number of traits related to dormancy and survival under stressful condition were affected by the removal of the *halR* gene. The four GO terms most significantly enriched for up-regulated genes in the exponential phase (cytoplasm, RNA binding, ATP binding, and tRNA binding) contained DEGs that could be generalised into three categories: DNA replication (helicases, gyrase, and topoisomerase), mismatch repair (*mutS*), and protein biosynthesis (rRNA and tRNA methyltransferases, aminoacyl-tRNA synthetases), corresponding to the enriched KEGG pathways (Aminoacyl-tRNA biosynthesis, DNA replication, Mismatch repair, and Protein export). Genes involved in metabolism of ribose, an important building block of DNA and RNA backbones, were amongst the genes with the highest FCs. This up-regulation occurring in the absence of HalR indicated that metabolism of nucleic acids was highly repressed in WT cells. Fatty acid biosynthesis was also amongst the up-regulated functions in both analyses, suggesting that the cell membrane biosynthesis was also repressed. Regulation of chromosomal replication and cell division by QS has been reported in *E. coli* (39, 40) and *P. aeruginosa* (41).

Genes involved in stress response and sugar uptake were negatively affected in the Δ*halR*::KanR mutant, which could be reflected in the down-regulation of two biological process terms (stress response and phosphoenolpyruvate-dependent sugar phosphotransferase system) and five pathways (phosphotransferase systems, carbon metabolism, microbial metabolism in diverse environments, fructose and mannose metabolism, and citrate cycle). Several genes involved in PTS with specificity towards mannitol and ascorbate, along with a number of other sugar permeases and transporters, such as maltose, fructose, and galactose, were amongst the genes with significant FC.

In the mid-exponential phase, the expression profile in the Δ*halR*::KanR mutant changed drastically. At this stage, many pathways involved in amino acid and fatty acid metabolism were down-regulated. In other words, these pathways were expressed at a higher level in the WT strain compared with the mutants, indicating that quorum sensing played a significant role in promoting metabolic activities in *H. alvei* FB1. The results of the GO enrichment analysis implied that phosphotransferase systems that were activated in WT earlier were now repressed, suggesting that these functions were more important in the early than in the mid-exponential phase. PTS are known to work not only as carbohydrate transport and modification systems in all phyla of bacteria, but also have regulatory roles in numerous cellular functions (42). In *E. coli*, glucose-specific PTS was found to inhibit other non-PTS sugar permeases influencing the availability of glucose *via* dephosphorylation of the EIIA subunit (43). This phenomenon, termed “catabolite repression”, contributes to the establishment of a hierarchy of sugar utilisation, which allows bacteria to utilise energy sources in an economical manner, as well as a regulatory circuit that acts according to nutrient availability (44).

The overall picture gained from these observations is that QS appears to be involved in repressing the functions involved in cell growth in an early growth phase, where the cells aree still in the process of recovery from stationary phase stress, in a typical ‘quorum-hindered apo-protein’ manner. The repressed genes may then be re-activated at a later stage when the DNA-bound apo-proteins are ‘hindered’ by the presence of AHL molecules at high cell density. Apo-HalR, on the other hand, could positively regulate the stress response and sugar uptake functions in the early exponential phase. PTS and various sugar transport systems that are active at low cell density could provide a mechanism for detection of nutrient availability and management of energy consumption. Whether these events of repression and activation are interdependent or under separate regulatory control will require further studies to resolve.

The observation that both mutants shared more than half of their enriched pathways in mid-exponential phase strongly suggests that QS regulation operated in a different manner different to the early exponential phase. In contrast with the expectation in a ‘quorum-hindered apo-protein’ model, very few DEGs showed contradicting expression level between the two mutants at this stage. Further study is required to provide a strong basis for the understanding of the regulatory mechanism of the HalR protein.

## Author statements

### Authors and contributors

JYT conceived and designed the experiments, performed the experiments, analysed the data, wrote the paper, and prepared figures and/ or tables. WSST conceived and designed the experiments, performed the experiments, and analysed the data. PC reviewed the manuscript. K-GC supervised the project, contributed reagents/ materials/ analysis tools. All authors approved the manuscript.

## Acknowledgements

K-GC thanks the research grants from the University of Malaya (FRGS grant FP022-2018A, PPP grant PG135-2016A, and HIR grant H-50001-A000027).

## References

1. Miller MB, Bassler BL. Quorum sensing in bacteria. Annu Rev Microbiol 2001; 55:165–199.

2. Parsek MR, Greenberg EP. Acyl-homoserine lactone quorum sensing in Gram-negative bacteria: a signaling mechanism involved in associations with higher organisms. Proc Natl Acad Sci U S A 2000; 97:8789–8793.

3. Duerkop BA, Ulrich RL, Greenberg EP. Octanoyl-homoserine lactone is the cognate signal for Burkholderia mallei BmaR1-BmaI1 quorum sensing. J Bacteriol 2007; 189:5034–5040.

4. Hao G, Burr TJ. Regulation of long-chain N-acyl-homoserine lactones in Agrobacterium vitis. J Bacteriol 2006; 188:2173–2183.

5. Pearson JP, Feldman M, Iglewski BH, Prince A. Pseudomonas aeruginosa cell-to-cell signaling is required for virulence in a model of acute pulmonary infection. Infect Immun 2000; 68:4331–4334.

6. Barnard AM, Bowden SD, Burr T, Coulthurst SJ, Monson RE, Salmond GP. Quorum sensing, virulence and secondary metabolite production in plant soft-rotting bacteria. Philos Trans R Soc Lond B Biol Sci 2007; 362:1165–1183.

7. Pierson LS, Gaffney T, Lam S, Gong F. Molecular analysis of genes encoding phenazine biosynthesis in the biological control bacterium, Pseudomonas aureofaciens 30-84. FEMS Microbiol Lett 1995; 134:299–307.

8. Rivas M, Seeger M, Jedlicki E, Holmes DS. Second acyl homoserine lactone production system in the extreme acidophile Acidithiobacillus ferrooxidans. Appl Environ Microbiol 2007; 73:3225–3231.

9. Tomlin KL, Malott RJ, Ramage G, Storey DG, Sokol PA, Ceri H. Quorum-sensing mutations affect attachment and stability of Burkholderia cenocepacia biofilms. Appl Environ Microbiol 2005; 71:5208–5218.

10. Labbate M, Queck SY, Koh KS, Rice SA, Givskov M, Kjelleberg S. Quorum sensing-controlled biofilm development in Serratia liquefaciens MG1. J Bacteriol 2004; 186:692–698.

11. Blana VA, Nychas GJ. Presence of quorum sensing signal molecules in minced beef stored under various temperature and packaging conditions. Int J Food Microbiol 2014; 173:1–8.

12. Bruhn JB, Christensen AB, Flodgaard LR, Nielsen KF, Larsen TO, et al. Presence of acylated homoserine lactones (AHLs) and AHL-producing bacteria in meat and potential role of AHL in spoilage of meat. Appl Environ Microbiol 2004; 70:4293–4302.

13. Christensen AB, Riedel K, Eberl L, Flodgaard LR, Molin S, et al. Quorum-sensing-directed protein expression in Serratia proteamaculans B5a. Microbiology 2003; 149:471–483.

14. Gram L, Christensen AB, Ravn L, Molin S, Givskov M. Production of acylated homoserine lactones by psychrotrophic members of the Enterobacteriaceae isolated from foods. Appl Environ Microbiol 1999; 65:3458–3463.

15. Ridell J, Korkeala H. Minimum growth temperatures of Hafnia alvei and other Enterobacteriaceae isolated from refrigerated meat determined with a temperature gradient incubator. Int J Food Microbiol 1997; 35:287–292.

16. Janda JM, Abbott SL. The genus Hafnia: from soup to nuts. Clin Microbiol Rev 2006; 19:12–18.

17. Milani C, Ticinesi A, Gerritsen J, Nouvenne A, Lugli GA, et al. Gut microbiota composition and Clostridium difficile infection in hospitalized elderly individuals: a metagenomic study. Sci Rep 2016; 6:25945.

18. Carding S, Verbeke K, Vipond DT, Corfe BM, Owen LJ. Dysbiosis of the gut microbiota in disease. Microb Ecol Health Dis 2015; 26:26191.

19. Jiang H, Ling Z, Zhang Y, Mao H, Ma Z, et al. Altered fecal microbiota composition in patients with major depressive disorder. Brain Behav Immun 2015; 48:186–194.

20. Tan JY, Yin WF, Chan KG. Quorum sensing activity of Hafnia alvei isolated from packed food. Sensors (Basel) 2014; 14:6788–6796.

21. Chin CS, Alexander DH, Marks P, Klammer AA, Drake J, et al. Nonhybrid, finished microbial genome assemblies from long-read SMRT sequencing data. Nat Methods 2013; 10:563–569.

22. Aziz RK, Bartels D, Best AA, DeJongh M, Disz T, et al. The RAST Server: rapid annotations using subsystems technology. BMC Genomics 2008; 9:75.

23. Aziz RK, Devoid S, Disz T, Edwards RA, Henry CS, et al. SEED servers: high-performance access to the SEED genomes, annotations, and metabolic models. PLoS One 2012; 7:e48053.

24. Angiuoli SV, Gussman A, Klimke W, Cochrane G, Field D, et al. Toward an online repository of Standard Operating Procedures (SOPs) for (meta)genomic annotation. OMICS 2008; 12:137–141.

25. Marchler-Bauer A, Bo Y, Han L, He J, Lanczycki CJ, et al. CDD/SPARCLE: functional classification of proteins via subfamily domain architectures. Nucleic Acids Res 2017; 45:D200–D203.

26. Galtier N, Gouy M, Gautier C. SEAVIEW and PHYLO_WIN: two graphic tools for sequence alignment and molecular phylogeny. Comput Appl Biosci 1996; 12:543–548.

27. Sharan SK, Thomason LC, Kuznetsov SG, Court DL. Recombineering: a homologous recombination-based method of genetic engineering. Nat Protoc 2009; 4:206–223.

28. Andrews S. FastQC: a quality control tool for high throughput sequence data. 2010. Available online at: http://www.bioinformatics.babraham.ac.uk/projects/fastqc

29. McClure R, Tjaden B, Genco C. dentification of sRNAs expressed by the human pathogen Neisseria gonorrhoeae under disparate growth conditions. Front Microbiol 2014; 5:456.

30. Tjaden B. De novo assembly of bacterial transcriptomes from RNA-seq data. Genome Biol 2015; 16:1.

31. Love MI, Huber W, Anders S. Moderated estimation of fold change and dispersion for RNA-seq data with DESeq2. Genome Biol 2014; 15:550.

32. Huang dW, Sherman BT, Lempicki RA. Bioinformatics enrichment tools: paths toward the comprehensive functional analysis of large gene lists. Nucleic Acids Res 2009; 37:1–13.

33. Huang dW, Sherman BT, Lempicki RA. Systematic and integrative analysis of large gene lists using DAVID bioinformatics resources. Nat Protoc 2009; 4:44–57.

34. Fuqua C, Greenberg EP. Listening in on bacteria: acyl-homoserine lactone signalling. Nat Rev Mol Cell Biol 2002; 3:685–695.

35. Subramoni S, Florez Salcedo DV, Suarez-Moreno ZR. A bioinformatic survey of distribution, conservation, and probable functions of LuxR solo regulators in bacteria. Front Cell Infect Microbiol 2015; 5:16.

36. Tsai CS, Winans SC. LuxR-type quorum-sensing regulators that are detached from common scents. Mol Microbiol 2010; 77:1072–1082.

37. Hao Y, Winans SC, Glick BR, Charles TC. Identification and characterization of new LuxR/LuxI-type quorum sensing systems from metagenomic libraries. Environ Microbiol 2010; 12:105–117.

38. Gray KM, Garey JR. The evolution of bacterial LuxI and LuxR quorum sensing regulators. Microbiology 2001; 147:2379–2387.

39. DeLisa MP, Wu CF, Wang L, Valdes JJ, Bentley WE. DNA microarray-based identification of genes controlled by autoinducer 2-stimulated quorum sensing in Escherichia coli. J Bacteriol 2001; 183:5239–5247.

40. Withers HL, Nordström K. Quorum-sensing acts at initiation of chromosomal replication in Escherichia coli. Proc Natl Acad Sci U S A 1998; 95:15694–15699.

41. Wagner VE, Bushnell D, Passador L, Brooks AI, Iglewski BH. Microarray analysis of Pseudomonas aeruginosa quorum-sensing regulons: effects of growth phase and environment. J Bacteriol 2003; 185:2080–2095.

42. Galinier A, Deutscher J. Sophisticated regulation of transcriptional factors by the bacterial phosphoenolpyruvate: sugar phosphotransferase system. J Mol Biol 2017; 429:773–789.

43. Kotrba P, Inui M, Yukawa H. Bacterial phosphotransferase system (PTS) in carbohydrate uptake and control of carbon metabolism. J Biosci Bioeng 2001; 92:502–517.

44. Brückner R, Titgemeyer F. Carbon catabolite repression in bacteria: choice of the carbon source and autoregulatory limitation of sugar utilization. FEMS Microbiol Lett 2002; 209:141–148.

